# Cancer cell-specific MHCII expression as a determinant of the immune infiltrate organization and function in the non-small cell lung cancer tumor microenvironment

**DOI:** 10.1101/2021.02.24.432729

**Authors:** Amber M. Johnson, Jennifer M. Boland, Julia Wrobel, Emily K. Klezcko, Mary Weisner-Evans, Lynn Heasley, Eric T. Clambey, Raphael A. Nemenoff, Erin L. Schenk

## Abstract

**Introduction:** In patients with non-small cell lung cancer (NSCLC), the prognostic significance of the tumor microenvironment (TME) immune composition has been demonstrated using single or dual-marker staining on sequential tissue sections. While these studies show that relative abundance and localization of immune cells are important parameters, deeper analyses of the NSCLC TME are necessary to refine the potential application of these findings to clinical care. Currently, the complex spatial relationships between cells of the NSCLC TME and potential drivers contributing to its immunologic composition remain unknown.

**Methods:** We employed multispectral quantitative imaging on the lung adenocarcinoma TME in 153 patients with resected tumors. On a single slide per patient, we evaluated the TME with markers for CD3, CD8, CD14, CD19, major histocompatibility complex II (MHCII), cytokeratin, and DAPI. Image analysis including tissue segmentation, phenotyping, and spatial localization were performed.

**Results:** Specimens where ≥5% of lung cancer cells expressed MHCII (MHCII^hi^ TME) demonstrated increased levels of CD4^+^ and CD8^+^ T cell and CD14^+^ cell infiltration. In the MHCII^hi^ TME, the immune infiltrate was closer to cancer cells and expressed an activated phenotype. Morphologic image analysis revealed cancer cells in the MHCII^hi^ TME more frequently interfaced with CD4^+^ and CD8^+^ T cells. Patients with an MHCII^hi^ TME experienced improved overall survival (p=0.046).

**Conclusions:** Lung cancer cell-specific expression of MHCII associates with levels of immune cell infiltration, spatial localization, and activation status within the TME. This suggest cancer cell-specific expression of MHCII may represent a biomarker for the immune system’s recognition and activation against the tumor.

## Introduction

The tumor microenvironment (TME) is a complex interface between cancer cells, stroma, and infiltrating immune cells. Multiple preclinical models of cancer have demonstrated that these interactions shape the natural history of tumor progression including the early events of primary tumor formation and how rapidly metastases develop(Reviewed in [1]). In patients with lung cancer, multiple analyses have demonstrated the prognostic significance of the TME immune composition[2]. High levels of immune infiltration within hematoxylin and eosin stained sections were associated with improved patient outcomes in a cohort of 1,546 patients with resected non-small cell lung cancer (NSCLC)[3]. Even in patients with stage I disease, immunologic characteristics of the TME including the proportion of FoxP3^+^ cells to total T cell infiltration correlated with disease recurrence in 956 patients with resected lung adenocarcinoma[4]. In addition to the differences in the numbers of immune cells, the location of specific populations within the NSCLC TME influences patient outcomes. In a cohort of 797 patients with resected lung cancer, high numbers of CD8^+^ T cells in the stroma associated with improved overall survival[5]. While these studies and others have demonstrated that relative abundance and localization of immune cells within the NSCLC TME are important parameters, deeper analyses of the NSCLC TME are necessary to refine the potential application of these findings to clinical care[1]. Currently, the complex spatial relationships between cells of the NSCLC TME and potential drivers contributing to its immunologic composition remain unknown.

Previously, we have reported that cancer cell-specific expression of major histocompatibility complex II (MHCII) modulates the TME within orthotopic mouse models of lung cancer[6]. Downregulation of MHCII in CMT167, a murine lung adenocarcinoma that expresses MHCII *in vivo*, resulted in fewer CD4^+^ and CD8^+^ T cells compared to the parental line[6]. Conversely, upregulation of MHCII in LLC, a murine lung adenocarcinoma that does not express MHCII *in vivo*, led to increased infiltration of CD4^+^ and CD8^+^ T cells compared to the parental line[6]. *Ex-vivo* experiments demonstrated MHCII expressing cancer cells presented antigen to CD4^+^ T cells and resulted in a Th1 phenotype. Studies of NSCLC tissues have demonstrated tumor cell positivity for MHCII and that cancer cell expression of MHCII associates with increased CD4^+^ T cell TME infiltration[6, 7]. We hypothesized NSCLC cancer cell-specific expression of MHCII would be a determinant of abundance and spatial organization of the immune infiltration within the NSCLC TME. We tested our hypothesis in a cohort of 153 patients with resected NSCLC adenocarcinoma and employed multispectral imaging for a deeper analysis of multiple cell phenotypes and their relative spatial distributions within the NSCLC TME. Here, we report NSCLC cancer cell-specific expression of MHCII results in a more abundant immune infiltration with a significant increase in proximity and interface of both CD4^+^ and CD8^+^ T cells to cancer cells. Notably, patients with NSCLC cancer cell-specific expression of MHCII experience improved overall survival.

## Methods

### Patient Studies

Tumor specimens were collected through a protocol approved by the Mayo Clinic Institutional Review Board and obtained from the Mayo Clinic Lung Cancer Repository. Written informed consent was obtained from all patients in accordance with the Declaration of Helsinki. Patients were identified from the tissue repository that underwent curative surgical resection of lung adenocarcinoma between 2004 and 2007, who had not previously received cancer directed therapy, and had available residual tumor specimens. The electronic medical record was reviewed and pertinent clinical data including age at time of surgery, gender, smoking status, pack years, and last follow up date or date of death was extracted. Patients were staged based on surgical findings using the 7th edition of the AJCC TNM system for NSCLC[8]. Formalin-fixed paraffin-embedded (FFPE) tissue blocks were sectioned into 5μm slides by the Pathology Research Core (Mayo Clinic, Rochester, MN). Hematoxylin and eosin slides were reviewed by a thoracic pathologist (JMB) to verify the presence of tumor.

### Tissue multiplex immunohistochemistry

Slides from FFPE blocks were stained using Opal IHC Multiplex Assay according to manufacturer’s protocol (PerkinElmer) as previously described[9]. Briefly, slides were sequentially stained for CD19 (clone BT51E, dilution 1:50, Leica), CD3 (clone LN10, dilution 1:500, Leica), CD14 (clone SP192, dilution 1:100, Abcam), CD8 (clone C8/144B, dilution 1:100, Dako), HLA-DR + DP + DQ (MHCII) (clone CR3/43, dilution 1:250, Abcam), cytokeratin (polyclonal Z0622, dilution 1:250, Dako) and DAPI (Akoya Biosciences) at the Human Immune Monitoring Shared Resource at CU Anschutz Medical Campus. Whole slide scans were acquired using the 10x objective via the Vectra imaging system (Perkin Elmer, version 3.0). Three to five regions from each slide containing tumor and stroma were selected utilizing Phenochart (v1.0.12, Perkin Elmer) for high resolution multispectral acquisition on the Vectra system at 20X magnification. The images were analyzed with inForm software (v2.4.8, Akoya) to unmix adjacent fluorochromes, subtract autofluorescence (**Figure 1A**), segment the tissue into tumor and stroma regions (**Figure 1B**), segment the cells into nuclear, cytoplasmic, and membrane compartments (**Figure 1C**), and to phenotype the cells according to morphology and cell marker expression (**Figure 1D**).

**Figure 1:**
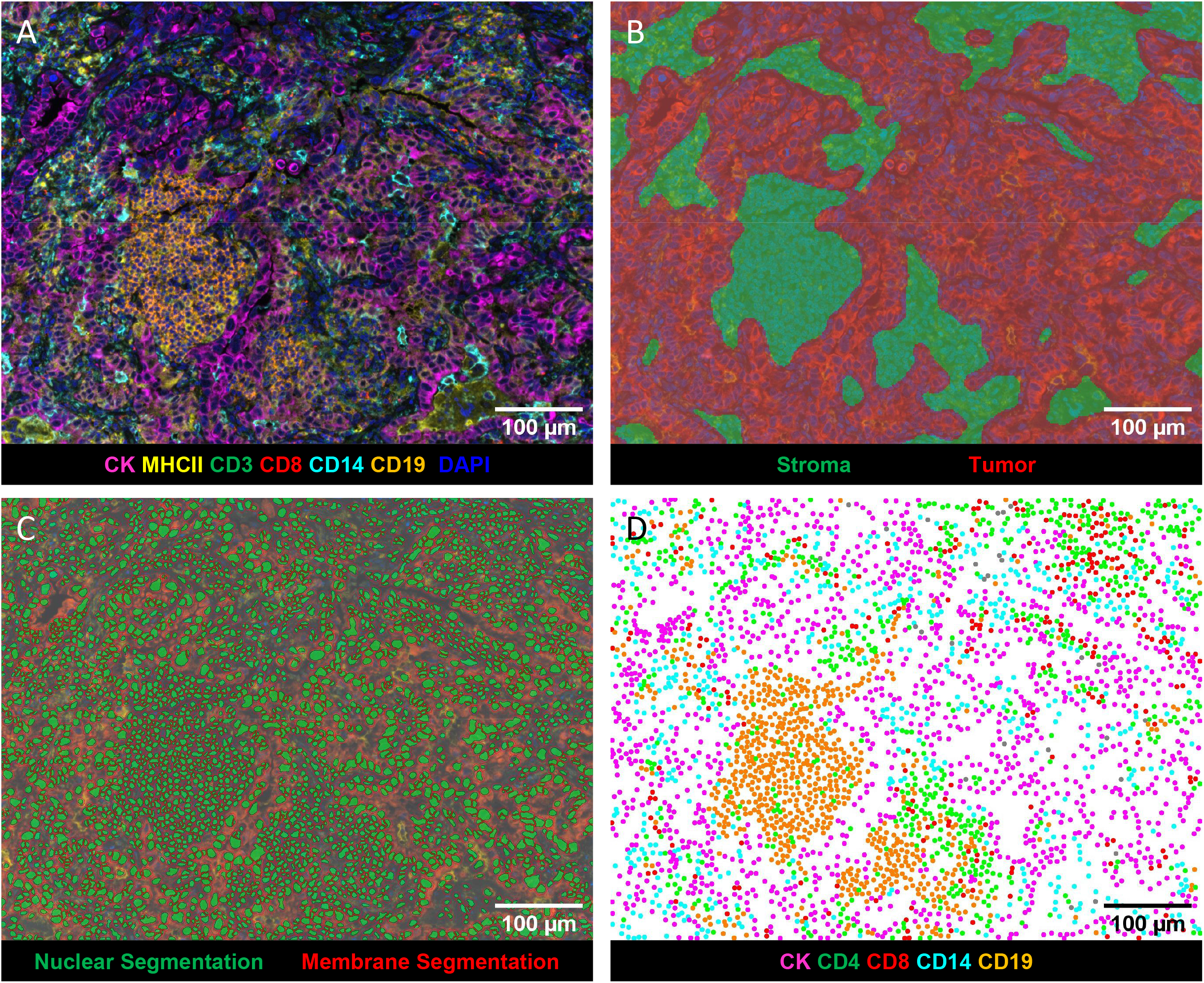
Representative images generated by inForm analysis. **A)** Spectrally unmixed composite image: pan-cytokeratin (CK) (magenta), CD3 (green), CD8 (red), CD14 (turquoise), CD19 (orange), MHCII (yellow), and DAPI (blue). **B)** Tissue segmentation map of tumor (red), and stroma (green) regions. **C)** Cell segmentation map: nucleus (green) and membrane (outlined in red). **D)** Phenotype map of CK^+^ cancer cells (magenta), CD4^+^ (CD3^+^CD8^−^) T cells (green), CD8^+^ T cells (red), CD14^+^ cells (turquoise), CD19^+^ B cells (orange).

### Data Analysis

inForm output data was analyzed using the R package Akoya Biosciences phenoptrReports to aggregate phenotype counts using count_phenotypes function for each slide and tissue category[10]. Nearest neighbor analysis was performed with the phenoptrReports nearest_neighbor_summary function[10]. The count_touching_cells function from phenoptrReports uses morphological analysis of nuclear and membrane segmentation maps to find touching cells of paired phenotypes[10]. To determine MHCII expression on cancer cells and immune cells, a threshold of membrane MHCII (Opal 690) mean of .05 positivity was determined by DAPI and MHCII single stain controls.

### Statistics

All statistical analysis performed was done with R version 4.0.3, R studio, or GraphPad Prism (v9.0.0, GraphPad Software). P values of p<.05 were considered significant.

## Results

### Cancer cell-specific MHCII expression associates with increased levels of TME immune infiltration

We quantified immune cell infiltration and MHCII expression in surgical resection specimens of patients with lung adenocarcinoma obtained from the Mayo Clinic Lung Cancer Repository. Patients were selected for multispectral imaging if they had no previous history of lung cancer and did not receive cancer directed therapy prior to surgery. Of the 153 patients evaluated 52.9% were women. Most of the patients were between ages 65-75 (75.8%), had a significant smoking history of 10 pack/years or more (73.2%), and were predominantly stage I (62%), specifically IA (41.8%), based on pathologic staging (**Table 1**). Initial quantification of the total number of immune cells within our cohort demonstrated a marked diversity degree and type of immune infiltration within each resection specimen (**Supplemental Figure 1**).

**Table 1.** Patient characteristics.

|  | n = 153 | % |
| --- | --- | --- |
| **Sex** |  |  |
| Male | 72 | 47.1 |
| Female | 81 | 52.9 |
| **Age groups** |  |  |
| <65 | 50 | 32.7 |
| 65-74 | 58 | 37.9 |
| ≥ 75 | 45 | 29.4 |
| **Smoking History** |  |  |
| Never | 22 | 14.4 |
| Light Smoker | 10 | 6.5 |
| Heavy Smoker | 112 | 73.2 |
| NA | 9 | 5.9 |
| **Stage (Pathological)** |  |  |
| IA | 64 | 41.8 |
| IB | 32 | 20.9 |
| IIA | 16 | 10.5 |
| IIB | 13 | 8.5 |
| IIIA | 21 | 13.7 |
| IIIB | 2 | 1.3 |
| IV | 5 | 3.3 |
| **Stage (Numeric)** |  |  |
| 1 | 96 | 62.7 |
| 2 | 29 | 19.0 |
| 3 | 23 | 15.0 |
| 4 | 5 | 3.3 |
| **Cancer Cell MHCII Status** |  |  |
| MHC II low < 5% | 56 | 36.6 |
| MHC II high ≥ 5% | 97 | 62.7 |

In our previous work, we reported that cancer cell-specific MHC class II expression alters the immune cell composition of the TME in murine models of lung cancer[6]. We therefore quantified the number of cancer cells positive for MHCII within each patient specimen as the percent of CK^+^ cells co-expressing MHCII (**Figure 2A**). To further determine the impact of cancer cell expression of MHCII on the TME, we classified tumor specimens as MHCII high or MHCII low based on previous work defining an MHCII high cancer cell population as ≥5% of tumor cells positive for MHCII[11, 12]. In our cohort, 63.4% (n = 97) of patients were found to have tumors with ≥5% of cancer cells expressing MHCII (MHCII^hi^) and 36.6% (n = 56) had tumors with <5% of cancer cells expressing MHCII (MHCII^lo^). Comparison of clinical characteristics between the MHCII^hi^ and MHCII^lo^ groups demonstrated no differences in patient distribution (**Table 2**). Differences were noted in the composition of the TME. The TME of MHCII^hi^ patients contained significantly increased total levels CD4^+^ and CD8^+^ T cell infiltrating the TME, and we also observed an increase in CD14^+^ cells compared to the TME of MHCII^lo^ patients (**Figure 2B and 2C**). Within the intratumoral compartment, MHCII^hi^ tumors contained more CD8^+^ T cells and increased CD14^+^ and CD19^+^ cells (**Supplemental Figure 2A**). Stromal infiltration of CD4^+^ T cells and CD14^+^ cells were higher in MHCII^hi^ tumors compared to MHCII^lo^ (**Supplemental Figure 2B**).

**Figure 2:**
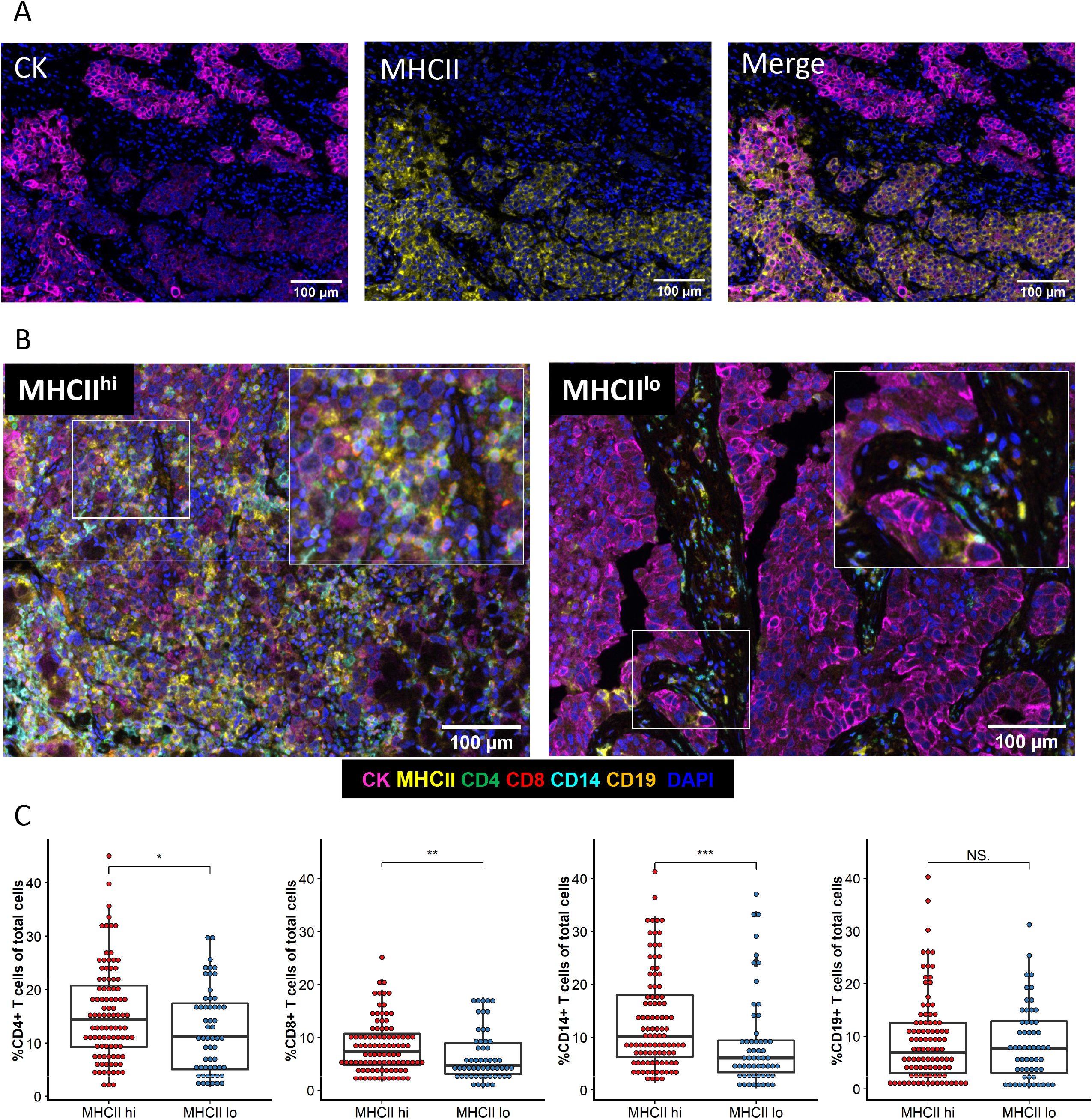
Immune infiltration is increased within the TME of MHCII^hi^ tumors. **A)** Representative images of single stain CK and MHCII and the merged image. **B)** Representative multispectral images of an MHCII^hi^ or an MHCII^lo^ TME. **C)** Percent of immune cells present within the TME of each patient by tumor MHCII status. Unpaired t-test. *p<.05, **p<.01, ***p<.001, NS not significant.

**Table 2.**
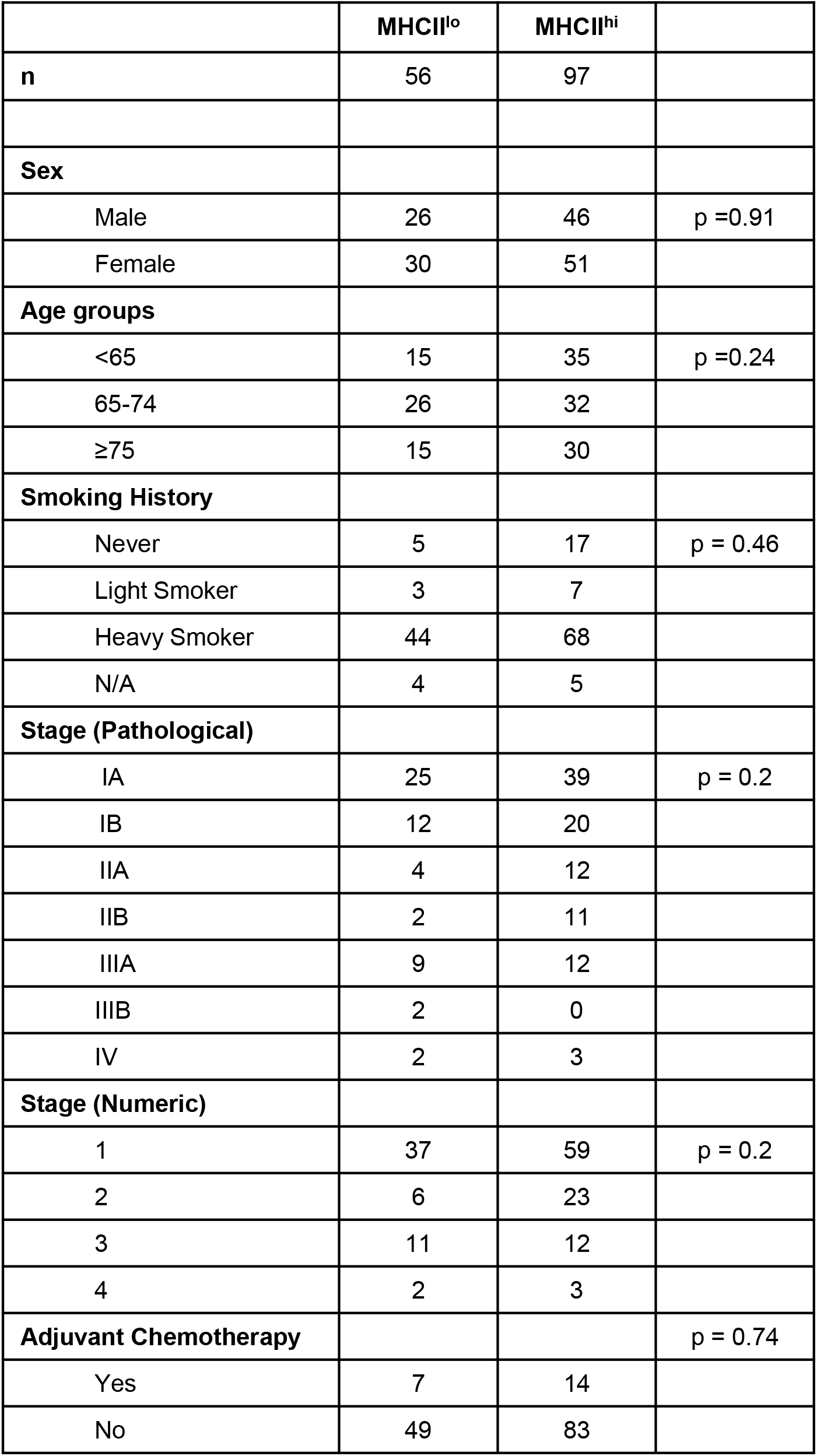
Patient characteristics by cancer cell MHCII status.

|  | MHCII<sup>lo</sup> | MHCII<sup>hi</sup> |  |
| --- | --- | --- | --- |
| **n** | 56 | 97 |  |
| **Sex** |  |  |  |
| Male | 26 | 46 | p = 0.91 |
| Female | 30 | 51 |  |
| **Age groups** |  |  |  |
| <65 | 15 | 35 | p = 0.24 |
| 65-74 | 26 | 32 |  |
| ≥75 | 15 | 30 |  |
| **Smoking History** |  |  |  |
| Never | 5 | 17 | p = 0.46 |
| Light Smoker | 3 | 7 |  |
| Heavy Smoker | 44 | 68 |  |
| N/A | 4 | 5 |  |
| **Stage (Pathological)** |  |  |  |
| IA | 25 | 39 | p = 0.2 |
| IB | 12 | 20 |  |
| IIA | 4 | 12 |  |
| IIB | 2 | 11 |  |
| IIIA | 9 | 12 |  |
| IIIB | 2 | 0 |  |
| IV | 2 | 3 |  |
| **Stage (Numeric)** |  |  |  |
| 1 | 37 | 59 | p = 0.2 |
| 2 | 6 | 23 |  |
| 3 | 11 | 12 |  |
| 4 | 2 | 3 |  |
| **Adjuvant Chemotherapy** |  |  | p = 0.74 |
| Yes | 7 | 14 |  |
| No | 49 | 83 |  |

### Increased spatial proximity between cancer cells and immune cells in the MHCII^hi^ TME

The proximity of immune cells to cancer cells within the TME is a critical determinant of immunologic activity, and we next determined the spatial localization of each cancer cell and immune cell by generating spatial coordinates for each cell type[1]. Distances between individual tumor cells and the nearest immune cell phenotypes were calculated. Spatial map views highlight the shorter distances between CK^+^ cancer cells and the nearest CD4^+^ T cell in an MHCII^hi^ compared to an MHCII^lo^ tumor (**Figure 3A**). Spatial analyses revealed an increased proximity of CD4^+^ and CD8^+^ T cells, CD19^+^, and CD14^+^ immune cells to the cancer cells in MHCII^hi^ tumors in contrast to MHCII^lo^ tumors (**Figure 3B**). In addition to the differences in cancer cell proximity, immune cells within the TME of MHCII^hi^ tumors more commonly expressed an activated phenotype compared to MHCII^lo^ tumors[13–15]. Higher levels of cell surface expression of MHCII on CD4^+^ and CD8^+^ T cells, CD14^+^ cells, and CD19^+^ cells were noted in MHCII^hi^ tumors compared to MHCII^lo^ tumors (**Supplemental Figure 3**). These differences remained significant when comparing tissue compartments reflecting an overall increased level in immune cell activity within the TME of MHCII^hi^ tumors (**Supplemental Figure 3**).

**Figure 3:**
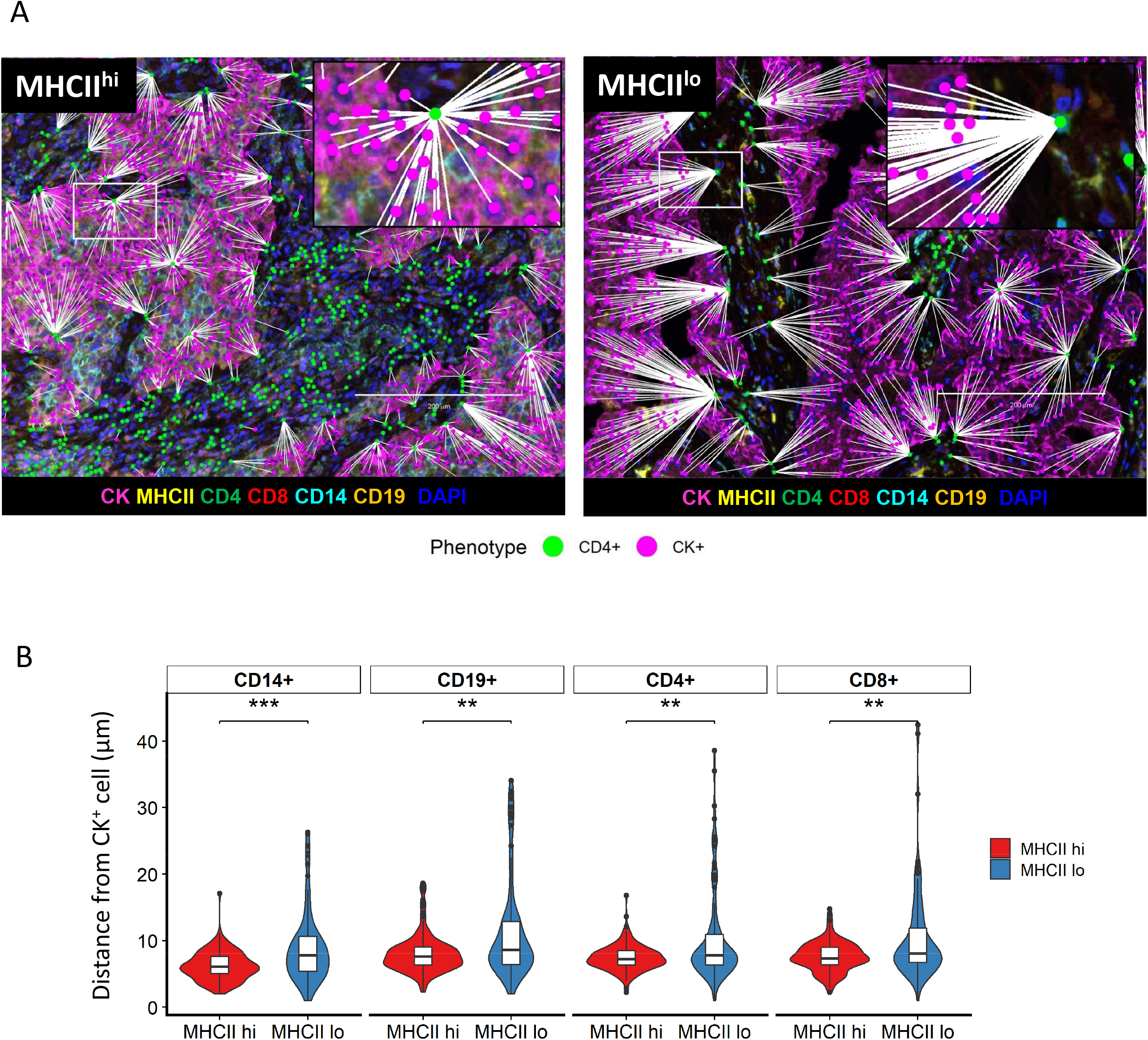
Increased spatial proximity of cancer cells to immune cells within the MHCII^hi^ TME. **A)** Representative spatial map image of the distance of nearest CD4^+^ cell (green) to CK^+^ cell (pink) in an MHCII^hi^ tumor or an MHCII^lo^ tumor. **B)** Average minimum distance of CK^+^ cells to CD14^+^, CD19^+^, CD4^+^, and CD8^+^ cells in MHCII^hi^ and MHCII^lo^ cohorts (n =153, Unpaired T-test, **p<.01, ***p<.001)

Ultimately, a successful anti-tumor response by the immune system culminates in CD8^+^ T cell recognition of the tumor and within our cohort we assessed this using morphologic analysis of the cell surface interface between pairs of cells[16]. Within the TME of MHCII^hi^ tumors, more cancer cells demonstrated a morphologic interface with CD8^+^ T cells in contrast to the TME of MHCII^lo^ tumors (**Figure 4A, Figure 4B**). CD8^+^ T cells within the TME of MHCII^hi^ tumors were more frequently interfaced with CD14^+^ cells, but not CD4^+^ T cells or CD19^+^ B cells (**Supplemental Figure 4A**). Similarly, tumor cells were more likely to share a surface interface with CD4^+^ T cells in MHCII^hi^ tumors (**Figure 4B, Figure 4C**). CD4^+^ T cells in the TME of MHCII^hi^ tumors were more frequently interfaced with CD14^+^ cells, but not CD8^+^ T cells or CD19^+^ B cells (**Supplemental Figure 4B**).

**Figure 4:**
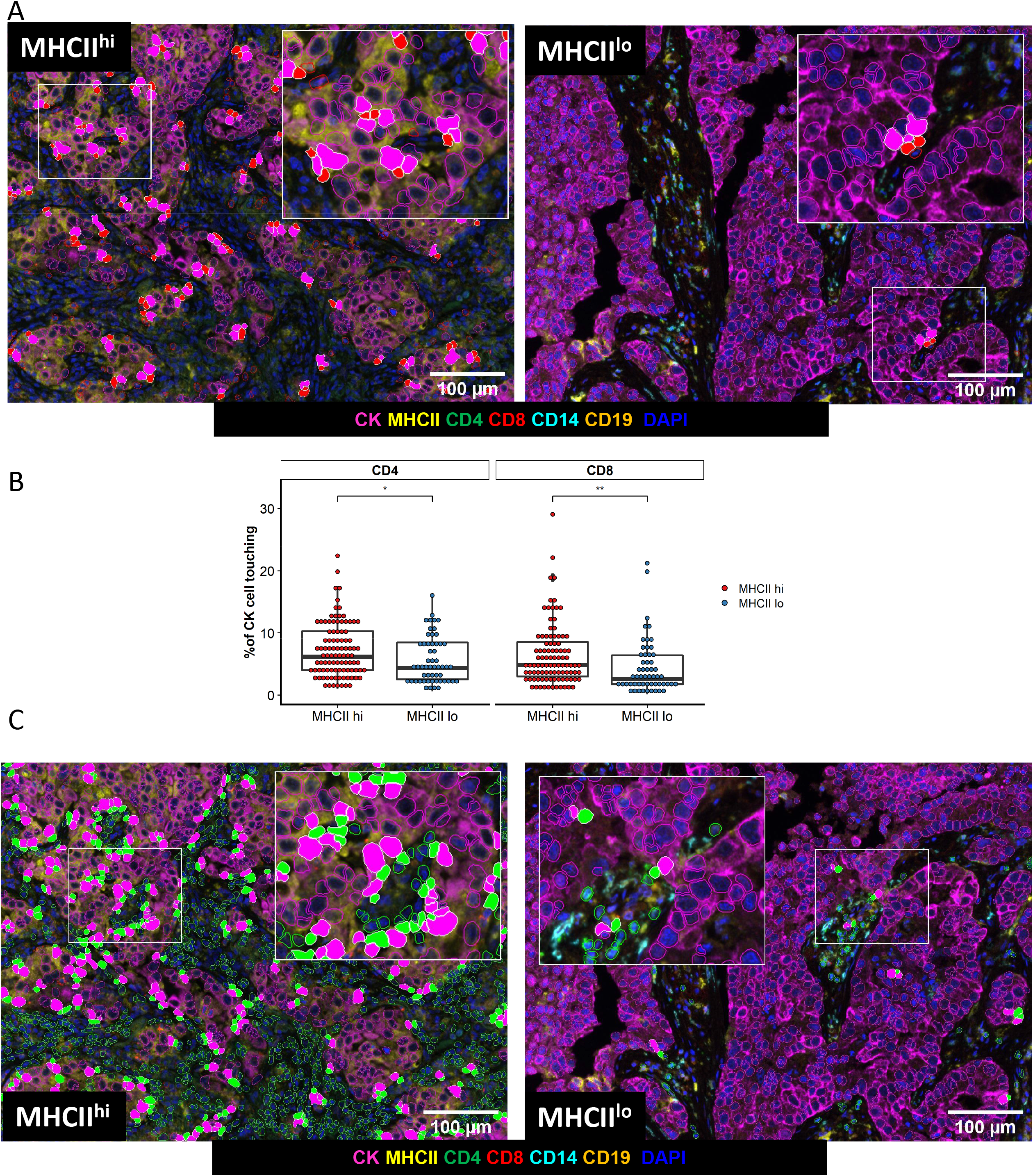
Cell-cell interface between cancer cells and T cells is more frequent in the MHCII^hi^ TME. **A)** Representative images of cell interface analysis between cancer cells and CD8^+^ T cells in an MHCII^hi^ and MHCII^lo^ tumor. **B)** Percent of cancer cells with a cell surface interface with CD4 ^+^ or CD8 ^+^ T cells based on tumor MHC status, unpaired t-test, *p<.05, **p<.01. **C)** Representative images of cell interface analysis between cancer cells and CD4^+^ T cells in an MHCII^hi^ or an MHCII^lo^ tumor.

### MHCII^hi^ Tumor Status Predicts Survival

As our analyses demonstrated MHCII^hi^ tumors have an increased quantity of total and activated immune cells and a higher frequency of T cell interfacing with cancer cells, we performed survival analyses on our patient cohort. Patients with MHCII^hi^ tumors experienced improved 5 year overall survival compared to patients with MHCII^lo^ tumors (**Figure 5A**). Improved survival was not observed when patients were grouped by high or low levels of immune cell infiltration based on the median of the entire cohort (**Figure 5B–5E**). In multivariate analysis, cancer cell-specific MHCII status was predictive of patient outcome (**Table 3**). Notably, levels of immune cell infiltration, grouped into high or low levels based on the entire cohort average, did not associate with 5 year overall survival.

**Figure 5:**
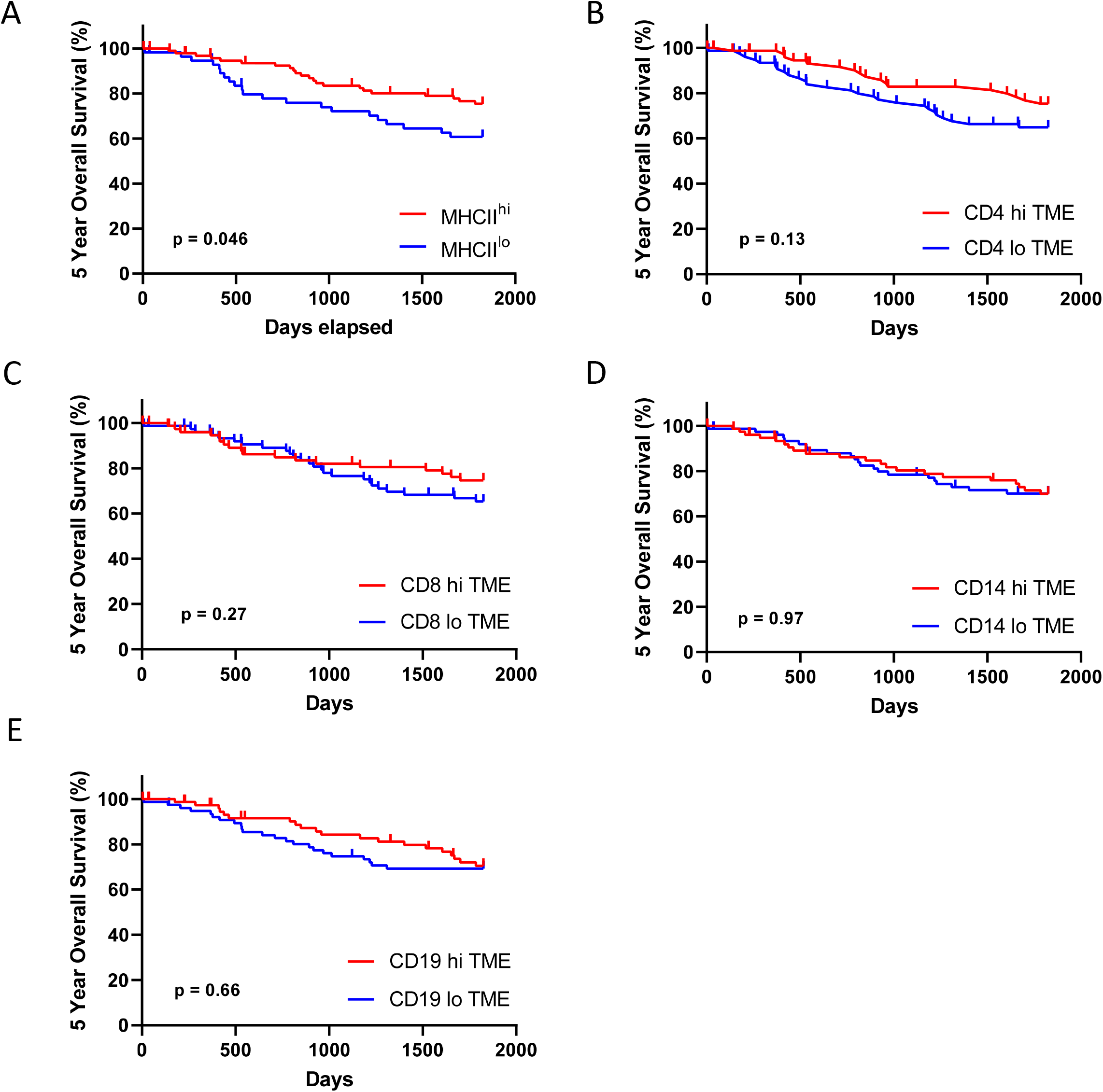
Five year overall. Stratified by **A)** Cancer cell-specific MHCII status, and levels of TME immune infiltration based on the cohort median of total **B)** CD4^+^, **C)** CD8^+^, **D)** CD14^+^, or **E)** CD19^+^ cells.

**Table 3.**
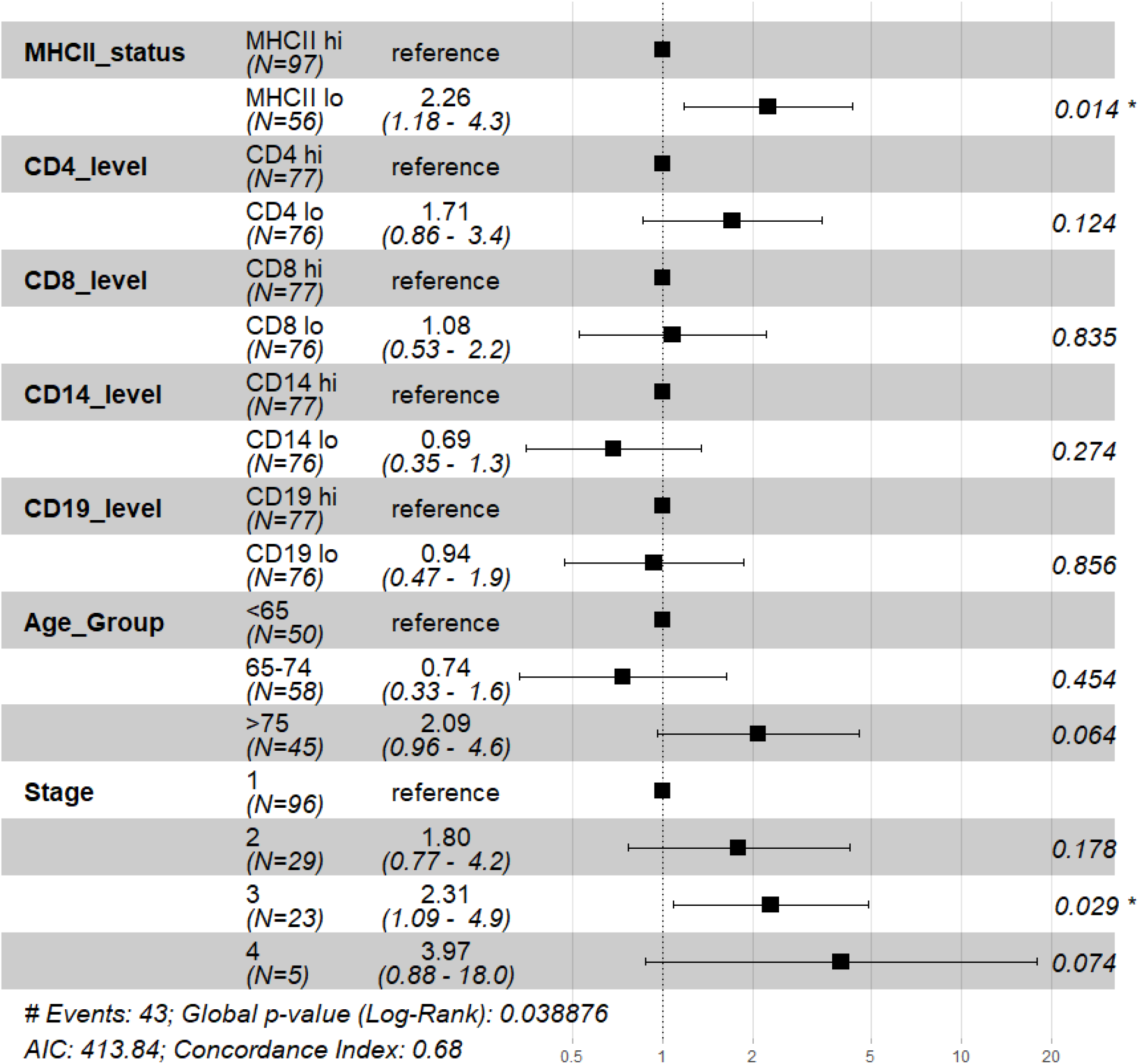
Multivariate analysis of 5 year overall survival.

## Discussion

While the composition of the TME has been recognized as a determinant of cancer progression and response to therapy, most analyses have focused on a limited number of cell types, and not examined multiple populations and their localization within the tumor. With the application of multispectral tissue imaging and bioinformatic approaches to analyze the TME of resected lung adenocarcinomas, we determined cancer cell MHCII expression associates with the organization and function of the immune infiltrate within the TME. The TME of MHCII^hi^ tumors contained higher levels of immune cell infiltration, similar to previous observations in melanoma, ovarian, and breast cancers[11, 17, 18]. In contrast to the previous studies of tumor MHCII expression and immune infiltration by IHC on serial sections, multiparameter tissue imaging allowed for simultaneous determination of multiple cell phenotypes and spatial coordinates in our cohort. With this level of single-cell resolution of phenotype and location, we found an increase in immune cell proximity to the tumor cells and degree of immune cell activation within the TME of MHCII^hi^ tumors. Importantly, within the TME of MHCII^hi^ tumors, cancer cells exhibited more cell surface in interfaces with CD4^+^ and CD8^+^ T cells, suggesting an increase in T cell recognition of cancer cells. In line with this hypothesis, we found patients with resected lung adenocarcinoma and an MHCII^hi^ tumor status experienced improved survival.

In part, our data of increased immune cell proximity and interface with cancer cells in MHCII^hi^ tumors strengthens the connection between improved patient outcomes and mechanistic data from animal models. In orthotopic models using the murine lung adenocarcinoma CMT-167 that expresses MHCII *in vivo*, knockdown of cancer cell-specific MHCII expression resulted in fewer activated CD4^+^ and CD8^+^ T cells within the lung cancer TME[6]. Additionally, knock down of MHCII expression in CMT-167 resulted in larger tumors that were less responsive to therapy[6]. Analogous findings were reported with the murine lung adenocarcinoma LLC, which does not express MHCII[6, 19]. Animals challenged with LLC cells transfected to express MHCII showed partial or complete abrogation of tumor growth, which was dependent on CD4^+^ and CD8^+^ T cells[19]. Notably, depletion of classical dendritic cells (cDC) in transgenic animals with the diphtheria toxin receptor under control of the CD11c promoter, did not impact the ability of animals to reject MHCII expressing LLC[19]. While other mononuclear phagocytes could serve as antigen presenting cells (APC) in the absence of cDC, one possibility is that the cancer cells expressing MHCII present antigen directly to the CD4^+^ T cells, resulting in activation. This in turn would lead to production of cytokines such as interferon-γ and augmentation of the immune response in a feed-forward manner [19]. This potential role of tumor cell as APC was further supported in an animal model of sarcoma where tumors were not eliminated unless the cancer cell expressed an MHCII neoantigen [20]. Our observation that cancer cells in MHCII^hi^ tumors more frequently interfaced with CD4^+^ T cells suggests this cell-cell interaction may play a similar role in immune mediated tumor control in patients with lung cancer.

For patients with clinical and radiographic early stage lung cancer, such as our cohort, the current standard of care is to first proceed with tumor resection[21]. Multiple ongoing clinical trials are evaluating whether neoadjuvant immune checkpoint inhibitors alone or in combination with chemotherapy improves patient outcomes versus standard of care (Reviewed in [22]). Our data associating tumor expression of MHCII with increased immune infiltration and proximity within the TME, indicators of immune recognition of the cancer cells, suggests tumor expression of MHCII may enrich for patients with lung cancer likely to respond to immunotherapy in this setting. Similar observations have been made in other malignancies where tumor expression of MHCII correlates with response to immunotherapy. Patients with metastatic melanoma who received PD-1 or PD-L1 immunotherapy experienced prolonged progression free and overall survival if ≥5% of tumor cells expressed MHCII[11]. Tumor cell expression of MHCII was also a key differentiator for progression free survival in patients with Hodgkin lymphoma who received PD-1 immunotherapy at least 1 year out from autologous stem cell therapy[23]. For patients with HER2 negative breast cancer, response to PD-1 immunotherapy associated with tumor expression of MHCII[24].

Our study provides translational rationale for evaluating cancer cell-specific expression of MHCII as a biomarker for immunotherapy responsiveness in patients with lung cancer. With our findings, several limitations exist that require the analyses of more modern cohorts. Initial studies can delineate whether cancer cell-specific MHCII associates with improved outcomes for patients with metastatic lung cancer receiving immunotherapy alone or in combination with chemotherapy in squamous and non-squamous histologies. Unlike the neoadjuvant setting, the use of immunotherapy in metastatic lung cancer for first line therapy has been standard of care for several years[21]. With this larger pool of potential patient specimens, further studies can be done with more complex multispectral tissue imaging in conjunction with tissue sequencing to better identify phenotypes and activation status of lymphocytes and leukocytes within the TME. Further work with modern patient cohorts could provide a more detailed insight into the role of cancer cell-specific MHCII and responsiveness to standard of care adjuvant chemotherapy than our cohort was able to address. An extension of our findings would be the hypothesis that patients with MHCII^hi^ NSCLC possess an active anti-tumor response and adjuvant chemotherapy would not improve survival, akin to observations in patients with deficient mismatch repair colorectal cancer[25]. Standard of care therapy for resected NSCLC continues to evolve and recently the presence of targetable driver oncogenes are now informing adjuvant therapy approaches[26]. While the role of the immune system in targeted therapy responsiveness is unknown, preclinical and translational data suggest engagement of the immune system promotes disease control[9, 27]. Whether cancer cell-specific expression of MHCII is a determinant of the TME and outcomes in patients with targetable oncogenes is unknown and remains an active area of investigation. In summary, we report that lung cancer expression of MHCII associates with levels of immune cell infiltration, spatial localization, and activation status within the TME. Along with pre-clinical data, our results suggest lung cancer expression of MHCII may serve as a biomarker for the immune system’s recognition and activation against the tumor. These insights may inform future approaches to predicting patient responsiveness to immunotherapy.

## Acknowledgements

ELS is supported by NIH grant K12 CA086913, ACS IRG #16-184-56 from the American Cancer Society to the University of Colorado Cancer Center, and a grant from the Cancer League of Colorado. RN is supported by NIH grant CA236222, DOD grant W81XWH1910221, and grants from the Cancer League of Colorado and the LUNGevity foundation.

The authors wish to thank the Pathology Research Core at Mayo Clinic for their assistance with specimen preparation. The authors wish to thank the Human Immunology Monitoring Shared Resource at the University of Colorado for their assistance with Vectra panel design and multispectral image acquisition, supported by the Cancer Center Support Grant (P30CA046934)

---

**Supplemental Figure 1:**
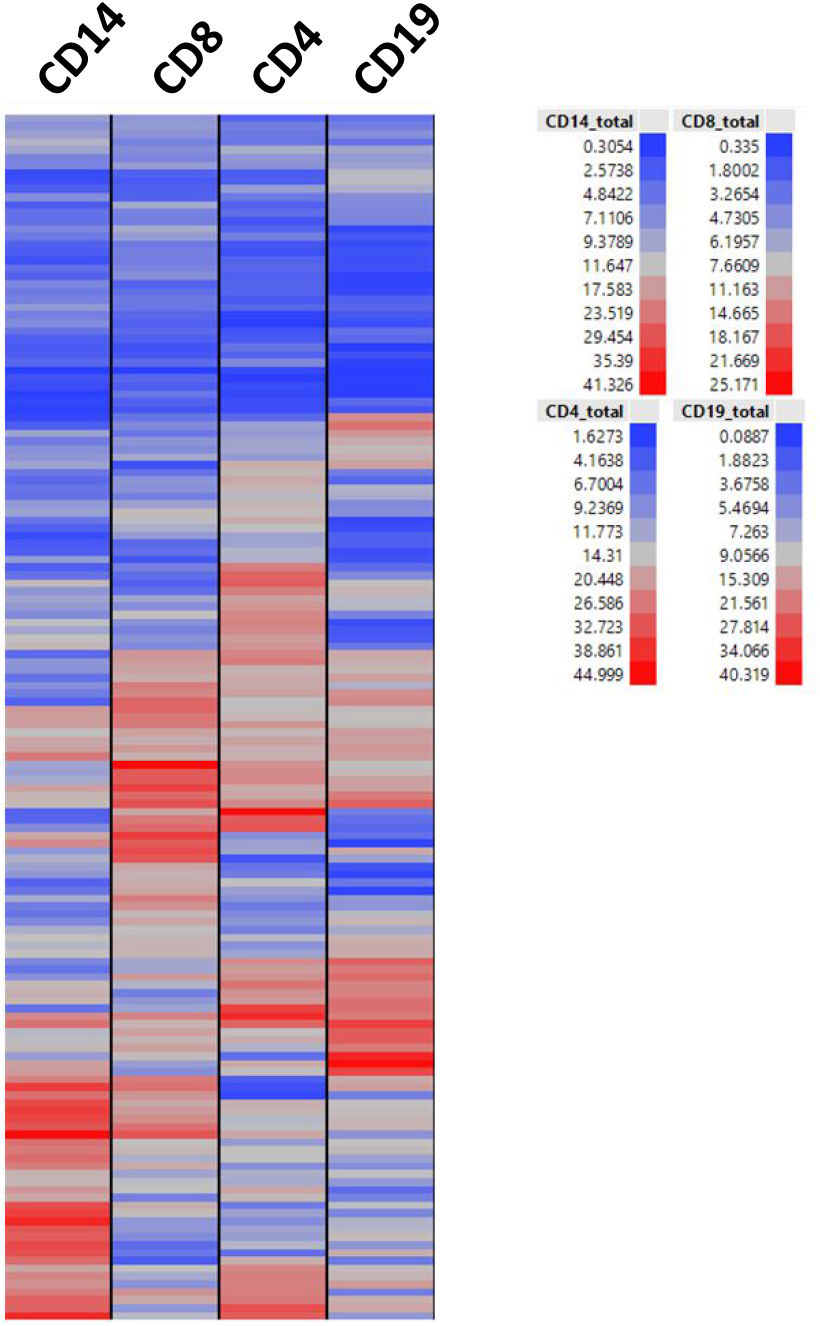
Immune infiltrate profiles of 153 lung adenocarcinoma specimens. Heat map rows represent individual patients and percent of each immune cell type out of total cells in the TME.

**Supplemental Figure 2:**
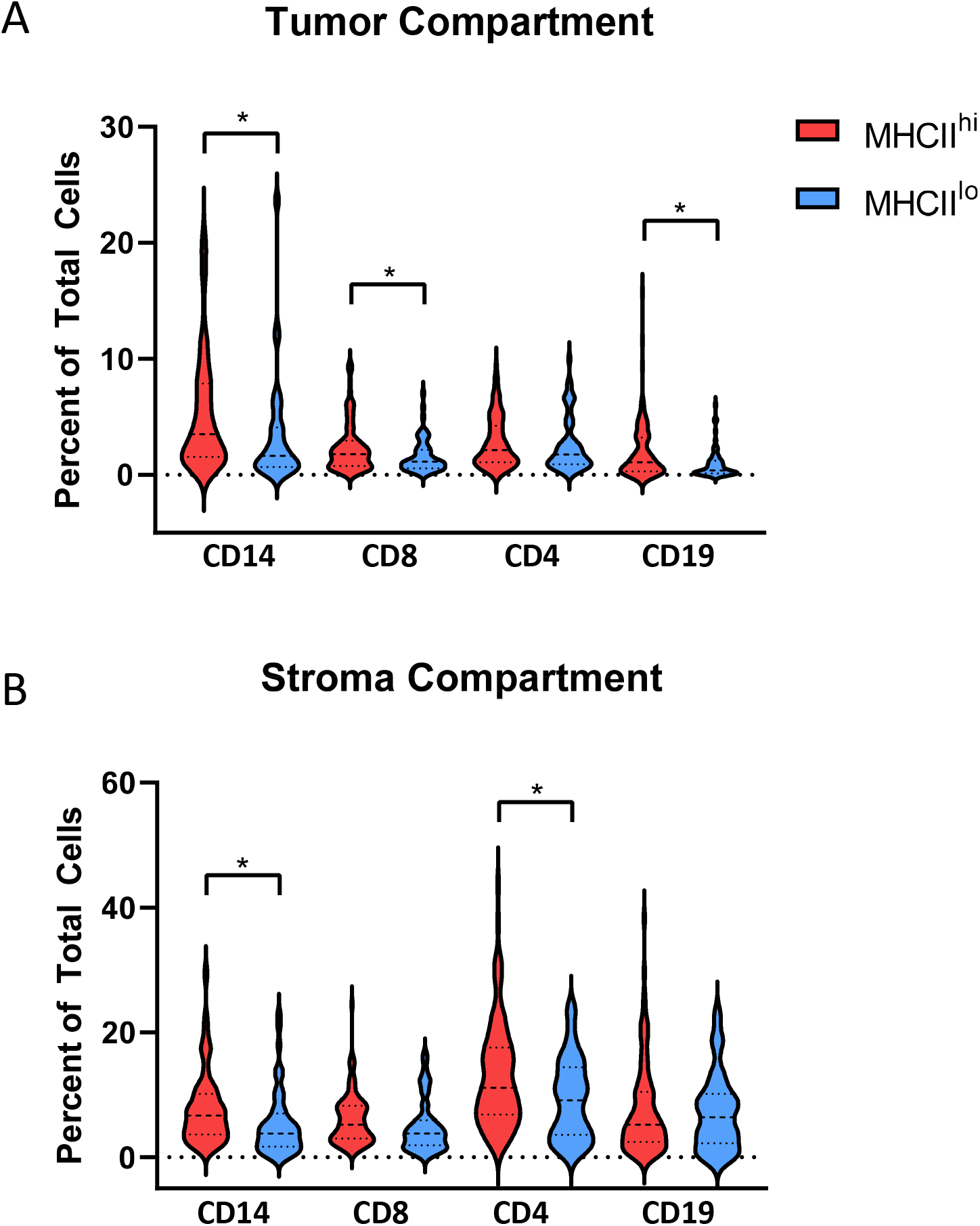
Proportion of Immune cells within the tissue compartments of the TME. **A)** Tumor, **B)** Stroma. *p<.05, unpaired t-test

**Supplemental Figure 3:**
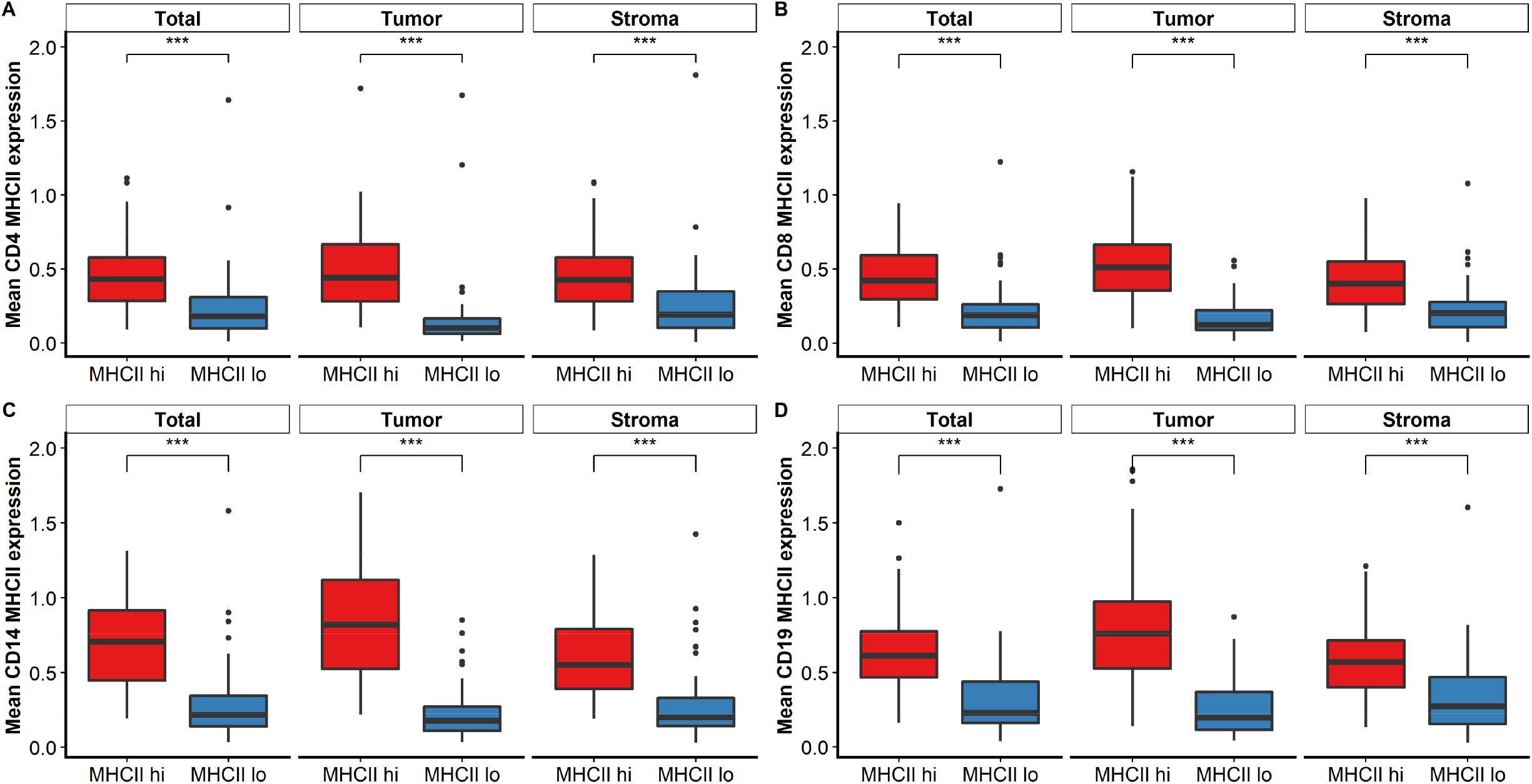
Immune cells within the MHCII^hi^ TME express higher levels of the activation marker MHCII. Differences in levels of immune cell expression of MHCII in the TME and stroma and tumor tissue compartments for **A)** CD4^+^, **B)** CD8^+^, **C)** CD14^+^, and **D)** CD19^+^ cells. Unpaired T-test, **p<.01, ***p<.001)

**Supplemental Figure 4:**
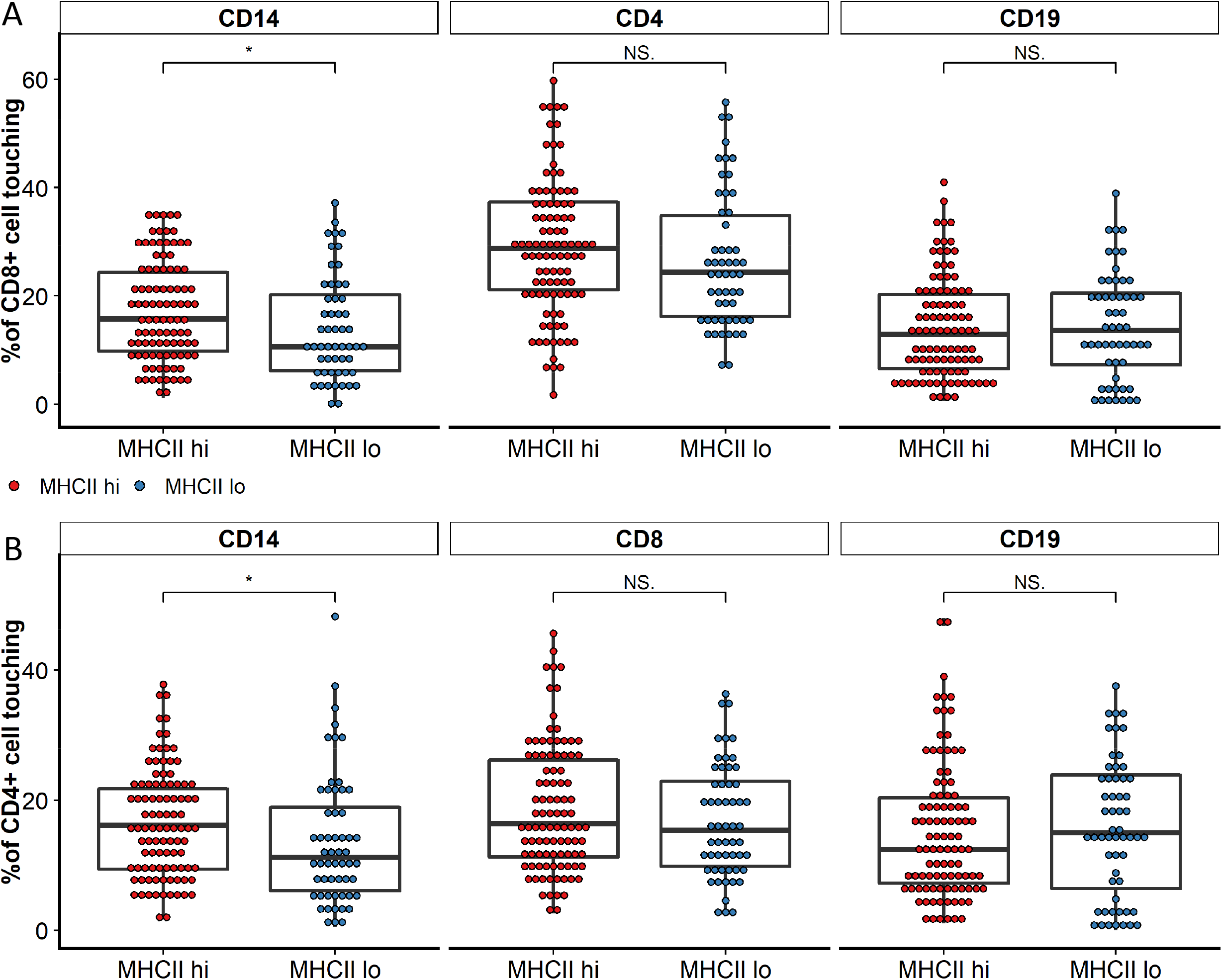
Cell-cell surface interface between T cells and other immune cells within the MHCII^hi^ or the MHCII^lo^ TME. Percent of **A)** CD8 T cells or **B)** CD4 T cells with morphologic interfaces with CD14, CD19, or other T cells in the MHCII^hi^ versus MHCII^lo^ TME. Unpaired t-test *p<.05

